# The limited effects of AAV2 vectors on host chromatin accessibility and nuclear architecture are consistent with a favorable safety profile

**DOI:** 10.1101/2025.09.09.675097

**Authors:** Joël W. Zöllig, Maija K. Pietilä, Kurt Tobler, Cornel Fraefel

## Abstract

Recombinant adeno-associated virus (AAV) vectors are widely used for gene therapeutics, yet their early effects on host chromatin remodeling and nuclear organization remain insufficiently characterized. In contrast, several DNA viruses are known to remodel host chromatin accessibility, for example HSV-1 in mammalian cells and baculoviruses in insect systems. Here we evaluated genome accessibility and nuclear organization of cultured primary human cells at 24 and 48 hours after infection with wildtype AAV2, single-stranded (ss) and self-complementary (sc) recombinant AAV2. Genome-wide ATAC-seq showed no detectable change in host chromatin accessibility at either time point. A DNase I digestion assay at five candidate loci supported this observation. In contrast, immunofluorescence imaging revealed modest decreases in histone H3, H3K27me3, and RNA polymerase II signals, consistent with reduced polymerase engagement under stress-linked pathways. H3K4me3 deviated from this pattern in G1 cells upon scAAV2 infection, where signals increased. Nuclear geometry shifted in parallel with protein signals, with changes in area, perimeter, convexity, and eccentricity. Markers of nuclear condensates also changed, including reduced fibrillarin, SP100, and SRSF2 intensities, altered object shape metrics, and higher counts of promyelocytic leukemia (PML) bodies and nuclear splicing speckles (NSs). Collectively, at the tested doses and times, AAV2-based vectors did not remodel chromatin accessibility at scale and induced only small changes in polymerase associated readouts and nuclear architecture. These data align with a favorable safety profile while highlighting assay limits and suggesting indirect stress related mechanisms.

## Introduction

Recombinant adeno-associated viruses (rAAVs) have emerged as safe and effective vectors for gene delivery in clinical applications, with several rAAV-based therapeutics already approved, including Luxturna for retinal dystrophy, Zolgensma for spinal muscular atrophy, and Hemgenix for hemophilia B (1–3). As of the time reported, AAV-based vectors had been evaluated in over 225 clinical trials addressing 55 distinct diseases, according to ClinicalTrials.gov data and related analyses (4). Ongoing studies are concentrated in five main therapeutic areas: blood disorders, central nervous system, eye disorders, lysosomal storage disorders, and neuromuscular disorders. AAV2 remains the most frequently used serotype, although the application of alternative serotypes has been steadily increasing (1,2).

AAVs are small, non-pathogenic viruses of the *Parvoviridae* family that replicate only in the presence of a helper virus. The wildtype AAV capsid contains a 4.7 kb single-stranded (ss)DNA genome in which inverted terminal repeats (ITRs) flank two major open reading frames, *rep* and *cap*, that in recombinant AAVs are replaced with a therapeutic sequence of interest, enabling the vector to deliver genetic material to target cells (5). Upon infection, rAAV genomes are maintained predominantly as extrachromosomal episomes, although integration into host chromosomal DNA can occur at low frequency (6–11). Meta analysis of rAAV-based clinical trials have shown that vector dosing spans a wide range, from approx. 10^9^ - 10^17^ vector genomes, with targeted delivery generally requiring lower doses than systemic administration (1).

In cell culture, induced pluripotent stem cells (iPSCs) and primary human fibroblasts have been reported to contain up to 100 and 5’000 rAAV vector copies per diploid genome when infecting at an MOI of 10’000 and 100’000, respectively (12). Human muscle cells can contain up to 160 vector copies per nucleus at three months after intramuscular administration of rAAV (13). Thus, rAAV-mediated gene therapy can introduce a significant amount of foreign DNA into the cell nucleus.

DNA viruses that replicate in the cell nucleus are known to drastically change the nucleus and chromatin organization. Assay for transposase-accessible chromatin using sequencing (ATAC-seq) shows that baculovirus infection remodels host chromatin and alters it more accessible in insect cells (14). Furthermore, besides chromatin reorganization, herpesviruses are known to impede several nuclear compartments, including promyelocytic leukemia (PML) nuclear bodies and nucleoli (15,16).

While AAV is a popular vector for human gene therapy, it remains unclear how the large numbers of rAAV genomes applied affect the nuclear architecture. The current study addressed this question using ATAC-seq and single-cell immunofluorescence imaging

## Materials and Methods

### Infection and transduction

1’350’000 BJ cells (ATCC CRL-2522) per 150 mm dish were used for ATAC-seq and qPCR. Immunofluorescence used 3,500 cells per well in glass bottom 96-well plates, cultured in Dulbecco’s Modified Eagle’s Medium (DMEM), supplemented with 10% fetal bovine serum (FBS). After 24 hours, cells were washed with serum-free DMEM and infected or transduced. Adsorption was 30 minutes at 4°C, then 60 minutes at 37°C. Unbound material was removed by washing with warm DMEM with 10% FBS, then cells were cultured for 48 hours.

### Nuclei isolation, DNA extraction and qPCR

At 24 or 48 hours, cells were harvested. Nuclei were isolated in hypotonic buffer (20mM Tris (pH 7.4), 10mM NaCl, 3mM MgCl_2_) supplemented with Nonidet-P40 at 0.5%, then vortexed for 10 seconds. DNA was extracted by phenol chloroform, precipitated with ethanol, resuspended in nuclease free water, and treated with RNase A.

Copy number of wtAAV2 and ssAAV2eGFP genomes were determined by SYBR Green qPCR using standard curves from the reference plasmids (Primers in Supplementary Tab. 1). Differences between groups were assessed with an unpaired two sample t-test.

### ATAC sequencing and sequence analysis

ATAC-seq libraries were prepared from 100’000 cells using the Active Motif ATAC-Seq Kit according to the manufacturer instructions. Libraries were quantified on an Agilent TapeStation and sequenced on an Illumina NovaSeq 6000 in paired end 2 x 150 bp mode. Raw reads were assessed with FastQ Screen (17) and FastQC (18), then aligned to the human reference genome GRCh38.p13 with Bowtie2 (19) within the SUSHI framework (20). Alignments were filtered, blacklisted loci were excluded (21), mitochondrial reads were removed, and a Tn5 offset was applied (22). Sorted and indexed BAM files were used for peak calling with MACS2 (23–25) in paired-end mode on the human genome. Quality control used FastQC and MultiQC (26). Differential accessibility was performed in R with DiffBind (27) on MACS2 peak sets, using consensus peaks, read counting, and contrasts by condition, with significance at false discovery rate less than 0.05 (28,29).

### Immunofluorescence and microscopy

Primary and secondary antibodies were applied in intercept buffer (Supplementary Tab. 3), with PBS washes between steps. Nuclei were stained with DAPI. For anti-H3 or anti-PCNA staining, a second permeabilization with 0.1% SDS was done. Imaging was performed on an ImageXpress Confocal HT.ai spinning disk confocal. Figures were prepared in napari, ImageJ and Affinity Designer (30,31).

### Segmentation, feature extraction and classification

Microscopy preprocessing, segmentation, and single cell quantification were performed in TissueMAPS (https://github.com/pelkmanslab/TissueMAPS). Nuclei were segmented from DAPI by Otsu thresholding with hole filling and declumping, small and large artifacts were removed. Intensity, texture, and morphology features were extracted. Random forest classifiers in TissueMAPS were trained iteratively as in CellClassifier (32). Cell cycle assignment followed published procedures using DAPI features, with k-means clustering in R (33,34).

### Data clean-up and normalization

Border touching objects and mitotic cells were excluded, and outliers were removed. Two independent experiments with three technical replicates were performed, and features were standardized within experiment relative to mock (i.e., not infected) by z-scoring to correct plate effects.

### LLPS compartment analysis using CellProfiler

Images were background normalized, nuclei were segmented by Otsu thresholding of DAPI. Nucleoli were segmented from fibrillarin, PML-bodies from SP100, and NSs from SRSF2. eGFP intensity was used to classify cells for eGFP-expressing or mock. For each segmented object, channel intensities and morphological features were quantified.

### DNase I digestion assay and qPCR

Protocol as described by Nepon-Sixt and Alexandrow (2019) (35), with DNase I treatment using 30 U for 10 min per sample. DNA was purified by phenol/chloroform extraction followed by ethanol precipitation, analysis by SYBR Green qPCR (Primers in Supplementary Tab. 3).

## Results

### Chromatin accessibility after AAV2 infection remains stable genome-wide and at selected loci

To determine whether AAV2 entry remodels human chromatin we performed an assay for transposase-accessible chromatin with high-throughput sequencing (ATAC-seq) at 24 and 48 hours post infection (hpi). Cells were infected with AAV2 (wildtype genome, wt) at multiplicity of infection (MOI) of 2’000 or 20’000 genome containing particles (gcp) per cell, and with single stranded (ss) or self-complementary (sc) AAV2 vectors carrying eGFP as transgene (ssAAV2 or scAAV2) at MOI of 20’000 gcp/cell. Genome-wide occupancy profiles across chromosomes 1 through X and Y in human primary fibroblasts are highly similar across all conditions and time points. No human genomic region showed significant changes in accessibility at either time point for any MOI (Fig. 1a). For wtAAV2, accessibility of the viral genome was modestly lower at 48 hpi compared to 24 hpi (Fig. 1b). Pairwise scatter plots histograms and Pearson correlation coefficients for eight samples defined by vector type, MOI, and time point yielded correlations above 0.80 in all comparisons and above 0.95 for ss and sc AAV2 at high MOI and late time point (Fig. 1c). This confirms that the main difference between the samples is due to the time difference, which explains 49% of variance (Fig. 1d).

**Fig. 1:**
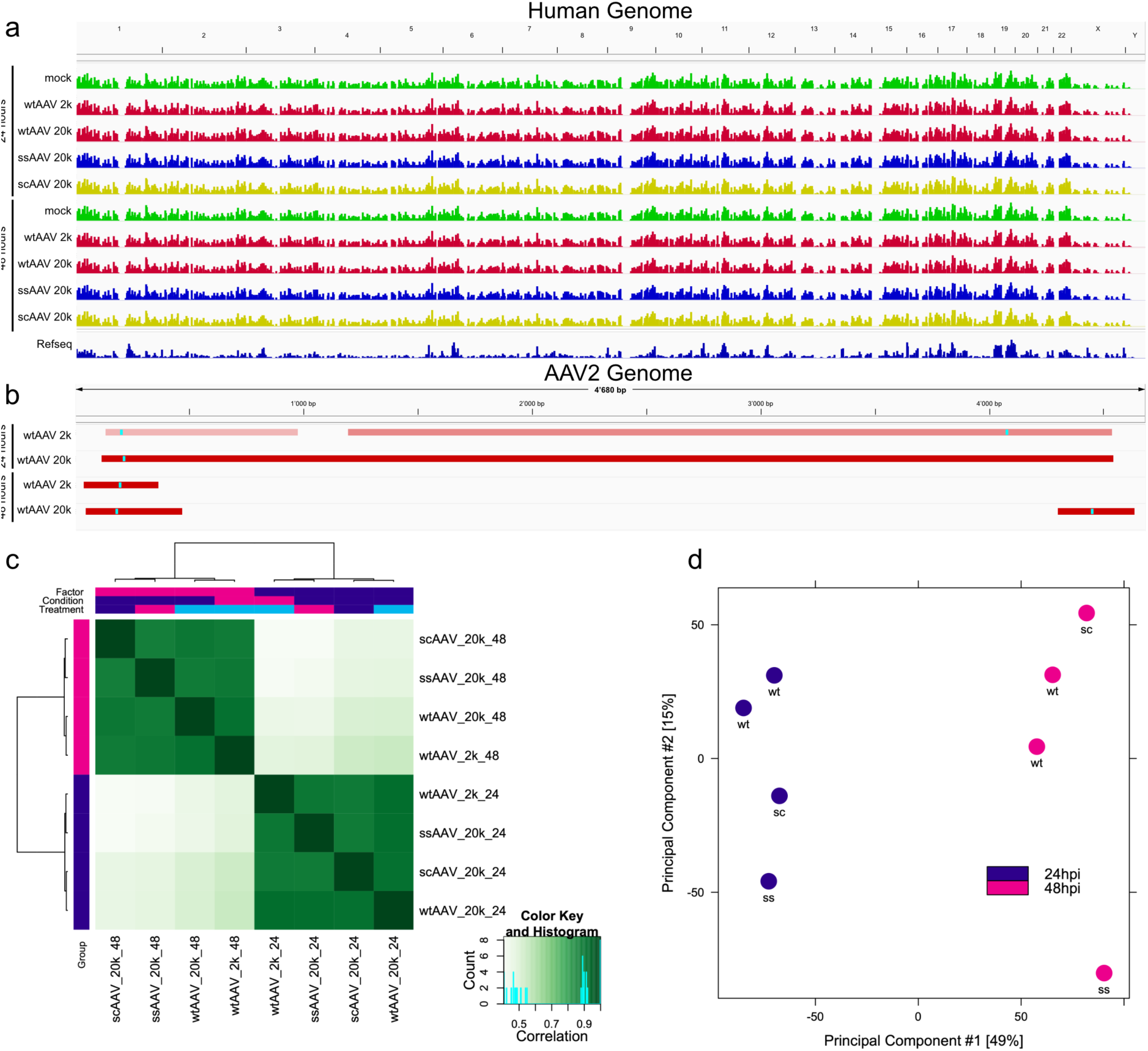
Genome-wide chromatin accessibility dynamics and correlation of AAV2 samples. (a) Chromosome occupancy of the human genome (hg38) at 24 and 48 hours after infection with wtAAV2, or eGFP encoding recombinant AAV2 vectors (ssAAV and scAAV) at MOI 2’000 or 20’000 gcp/cell (2k, 20k). Peaks demonstrate transposable-accessible (i.e., open), chromatin across chromosomes 1 through XY in primary human fibroblasts. (b) Chromosome occupancy for AAV2 alone after 24 and 48 hours at MOI of 2’000 or 20’000 gcp/cell (2k, 20k). Both IGV-snapshots in (a) and (b) show representative data. (c) Pairwise comparison and clustering of eight samples defined by vector type (wtAAV, ssAAV, scAAV), MOI (2k, 20k), and harvest time (24 or 48 hours). The lower triangle shows scatter plots of raw occupancy values, the diagonal shows sample histograms, the upper triangle shows Pearson correlation coefficients with color indicating strength. (d) Principal component analysis of the same eight samples as in (c).

### DNase I digestion assay at candidate genes confirms unchanged chromatin accessibility

To validate the global findings at candidate loci, we carried out a DNase I digestion assay. First, the extracted DNA was treated for 10 min either with DNase I, or digestion buffer without DNase I as control. The DNA was then subjected to quantitative (q)PCR with primer pairs targeting the tightly packaged alpha satellite DNA stretch D7Z1 for normalization, as well as the five genes ACTB (β-actin, cytoskeletal protein supporting cell structure and motility), GNG7 (G-protein gamma subunit, modulates GPCR signaling pathways), IFNB1 (type I interferon β, drives antiviral innate immune responses), OSBPL8 (lipid transfer protein, regulates cholesterol and phospholipid homeostasis), and PCNA (DNA sliding clamp, coordinates replication and repair with polymerases) (36). After expressing each target relative to D7Z1, chromatin openness was assessed by calculating the ΔΔCt, defined as the Ct difference of DNase I-treated samples relative to the corresponding non-DNase I-treated control. A ΔΔCt = 0 indicates that the chromatin is tightly packaged and therefore protected, whereas a ΔΔCt > 0 demonstrates the openness of the chromatin (Fig. 2). No statistical differences in relative accessibility across all conditions at both time points were found, thus confirming the findings from the ATAC-seq (Fig. 1). Representative IGV snapshots of the ATAC tracks at these loci show no detectable change in chromatin openness relative to mock (i.e., not infected).

**Fig. 2:**
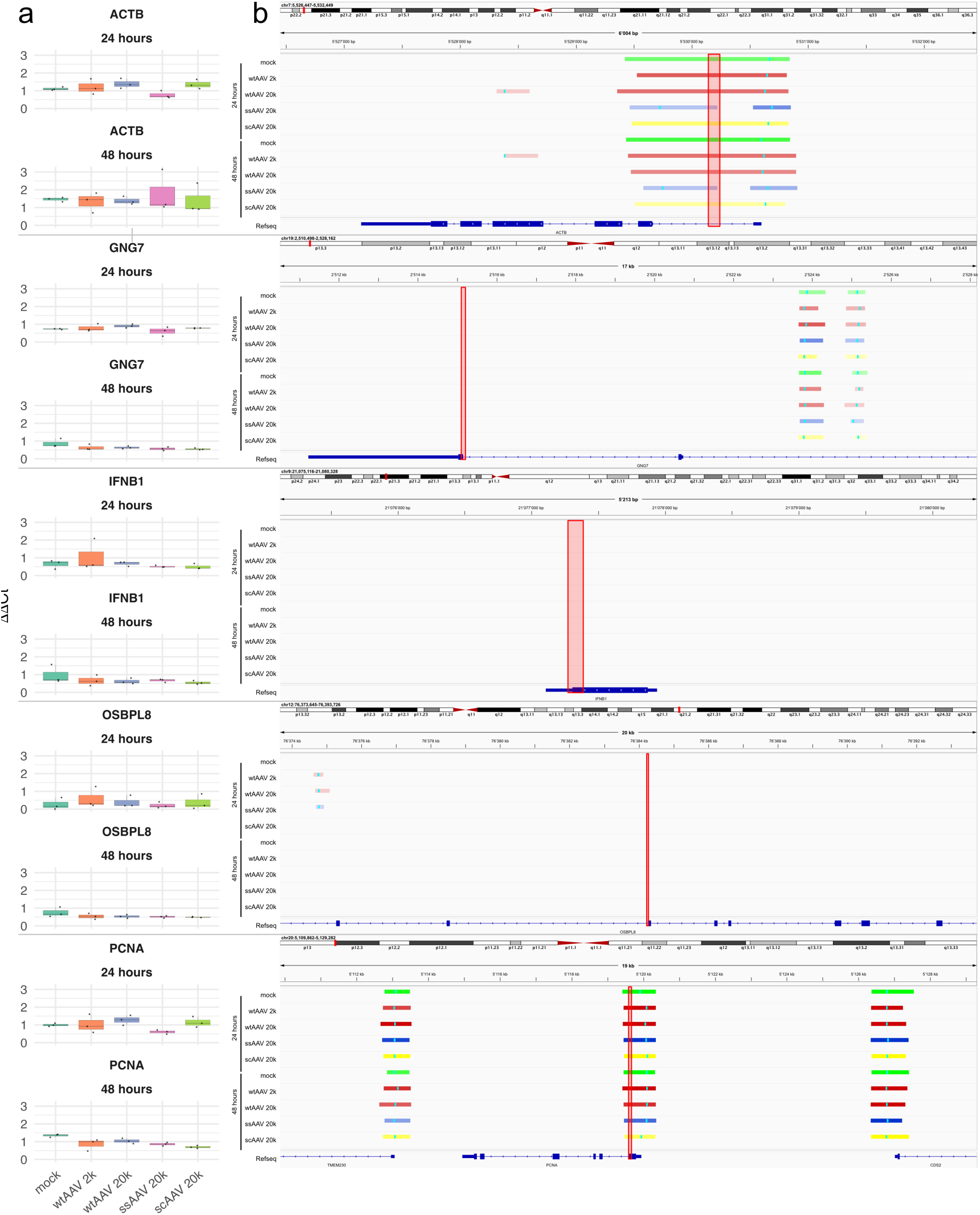
DNase I digestion and ATAC-seq confirm stable chromatin accessibility at targeted loci following infection with wtAAV2 or recombinant AAV2 (ssAAV2, scAAV2) (a) ΔΔCt from the DNase I digestion assay performed in biological triplicate for the genes ACTB, GNG7, IFNB1, OSBPL8, and PCNA after 24 and 48 hours of infection. Primary human fibroblasts were mock-infected, or infected with either wtAAV2 at MOI 2’000 or 20’000 gcp/cells (2k, 20k), or eGFP-encoding recombinant AAV2 vectors (ssAAV2, scAAV2) at MOI of 20’000 gcp/cell (20k). Raw Ct values were first normalized to D7Z1 as internal reference, then chromatin openness assessed by calculating the difference between DNase I-treated and non-treated samples, shown as ΔΔCt. ΔΔCt = 0 indicates packaged chromatin, ΔΔCt > 0 demonstrates open chromatin, the higher the value, the more open it is. The statistical analysis used one way ANOVA for most comparisons, and the Kruskal-Wallis test for IFNB1 and OSBPL8 at 24 hpi because normality was rejected. (b) Representative IGV snapshots of the ATAC sequencing covering the genes in (a), the reference gene, and the PCR amplification product (red square).

### AAV2 vectors transduce nuclei efficiently with modest effects on cell cycle

Next, primary human fibroblasts were infected with ssAAV2 or scAAV2 at MOI 20’000 gcp/cell, and eGFP-expression and localization were measured at 48 hpi by high-content screening immunofluorescence microscopy (HCS-IF) (Fig. 3a). Nucleus segmentation is shown in red, and eGFP signal indicates transgene expression. Total cell number per well was lower than mock for both vectors (ssAAV2 and scAAV2) (Fig. 3b). At 48 hours, the fraction of eGFP-expressing cells was equivalent between ssAAV2 and scAAV2 within a 5% margin (Fig. 3c). Cell cycle classification based on nuclear DAPI intensity and nuclear area (37) indicated a shift from G1 into S/G2 phase in vector-treated cells, while mock remained enriched in G1. Among eGFP-expressing cells, ssAAV2 showed a lower G1 fraction than scAAV2, indicating a shift toward S/G2-phase. The difference was significant but modest relative to the contrast with mock (Fig. 3d). Lastly, qPCR analysis of nuclear extracts revealed similar viral genome copy numbers between wtAAV2-infected and ssAAV2 vector-infected cells after 48 hours (Fig. 3e).

**Fig. 3:**
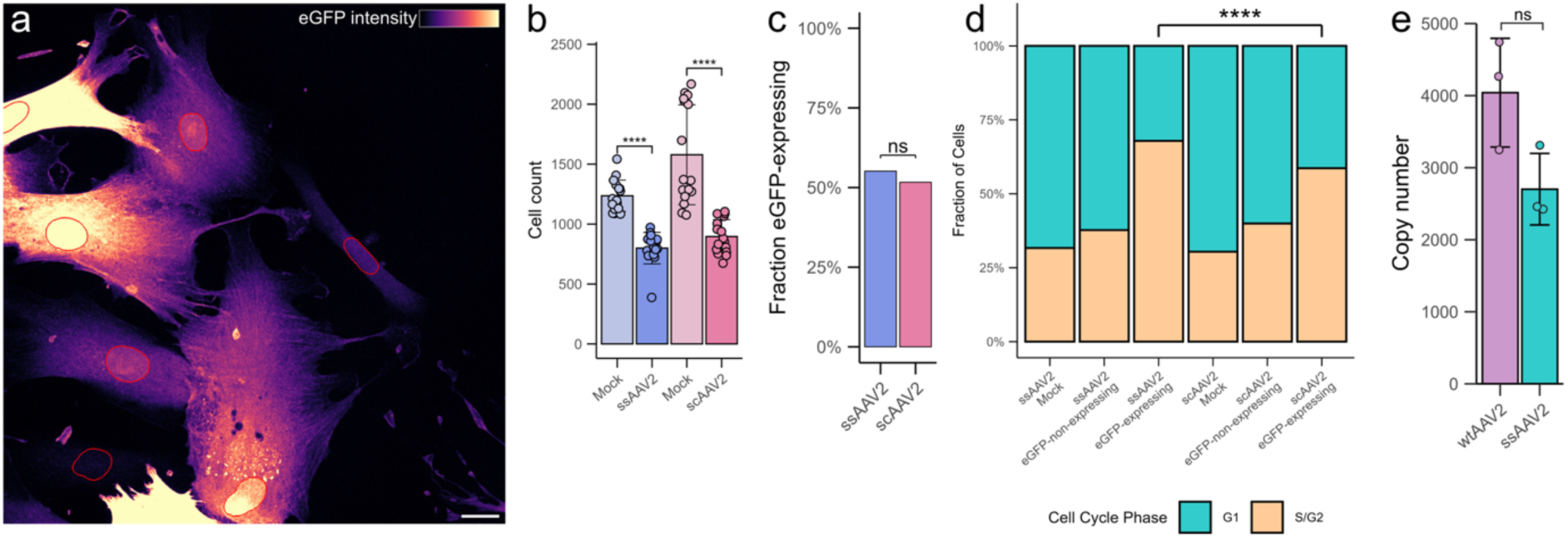
AAV2 at 48 hours increases eGFP, reduces cell number, shifts cells toward S or G2, and accumulates high nuclear genome copies. (a) Representative eGFP image of ssAAV2-infected primary human fibroblast cells at 48 hpi at MOI of 20’000 gcp/cell, nuclei outlined in red, scale bar 25 µm. (b) Cell counts for the infection of mock, ssAAV2, and scAAV2, at 48 hpi and MOI 20’000 gcp/cell (Wilcoxon rank sum tests, two condition comparisons). (c) Fraction of eGFP expressing cells at 48 hpi for ssAAV2 and scAAV2, equivalence tested on the absolute difference with a 5% margin using the two one-sided tests procedure at α = 0.05. The 90% confidence interval for ssAAV2 minus scAAV2 was 2.5% to 4.5%, which lies within the margin. (d) Cell cycle phase distribution determined from DAPI mean intensity and nuclear size using k-means clustering. Differences in cell cycle fractions were tested with a binomial generalized linear model on G1 versus S/G2 counts, with estimated marginal means and Tukey adjusted pairwise contrasts. (e) Number of genome copies in primary human fibroblasts upon infection with AAV2 or ssAAV2 at MOI of 20’000 gcp/cell at 48 hpi, determined by qPCR under the assumption of all genomes being single-stranded (unpaired two-sample t-test). * p < 0.05, ** p < 0.01, *** p < 0.001, **** p < 0.0001, ns = not significant.

### ssAAV2 and scAAV2 modestly reduce histone and RNAP II signals

Next, we profiled chromatin markers and RNA polymerase II by HCS-IF at MOI 20’000 gcp/cell. Cells were grouped by eGFP-expression and by cell cycle state, G1 or S/G2. All readouts were z-score normalized and the Δz relative to mock calculated (Fig. 4a). Most nuclear mean intensities for histones H3, H3K27me3, and the RNA polymerase II (RNAP II) markers show a reduction relative to mock, consistent with a mild reduction in polymerase engagement and chromatin marker abundance. However, H3K4me3 behaves differently. While many conditions show reduction, a subset shows maintained or increased signal, including a significant rise in cells which are in G1-phase and eGFP-expressing upon scAAV2 infection.

**Fig. 4:**
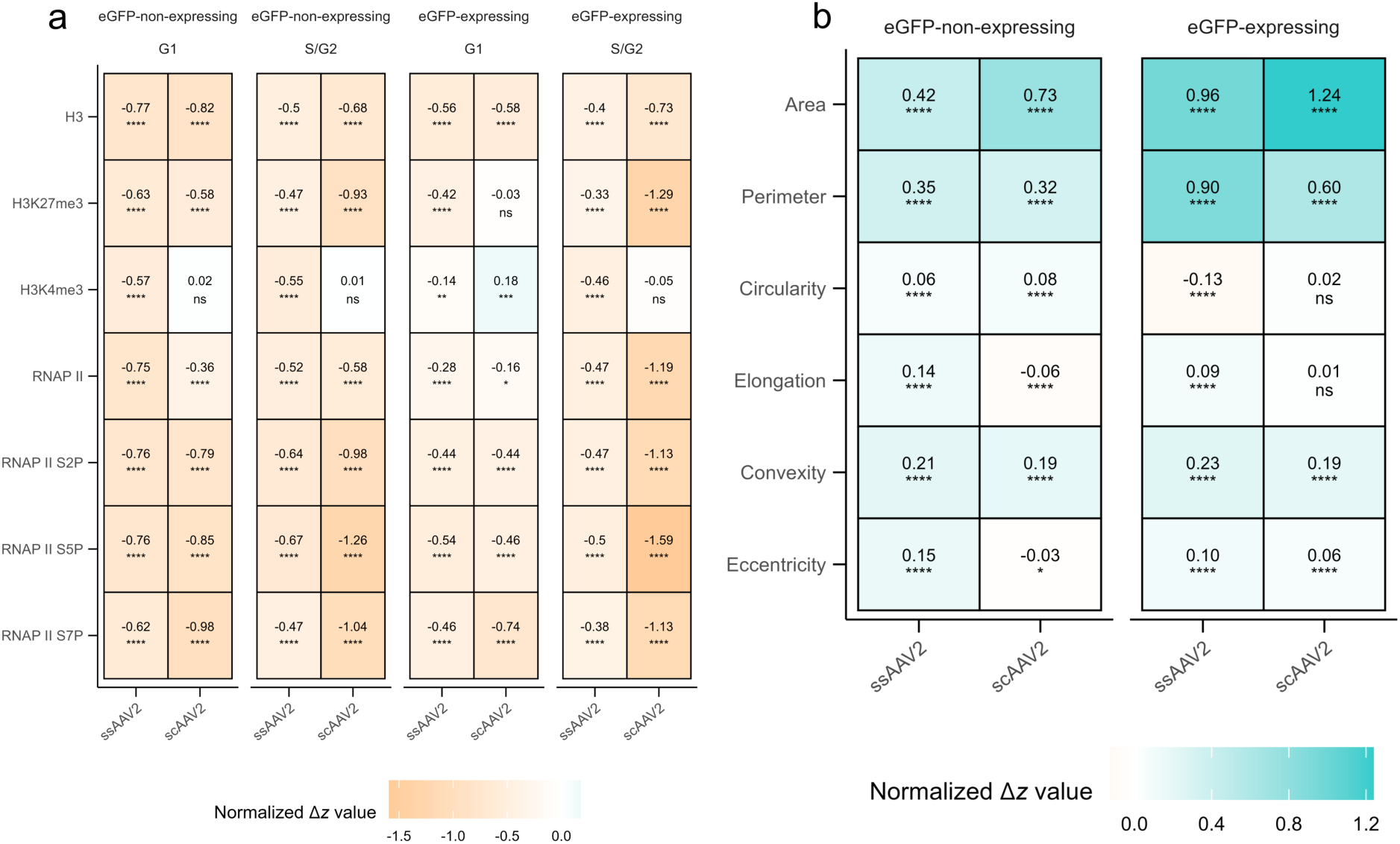
Chromatin markers, RNA polymerase II, and nuclear morphology upon ssAAV2 and scAAV2 infection. (a) Heatmap of Δz for mean nuclear intensities of histones H3, H3K27me3, H3K4me3, RNA polymerase II (RNAP II), and its phosphorylation RNAP II S2P, S5P, S7P, grouped by eGFP-expressing and eGFP-non-expressing cells, divided into G1 and S/G2 subgroups, comparing ssAAV2 and scAAV2 to mock. Values were z-score normalized, then Δz was calculated relative to mock. Cells were classified into G1 and S/G2 based on their DAPI-intensity and nuclear area. Asterisks indicate significant change relative to mock. (b) Heatmap of Δz for nuclear morphology features area, perimeter, circularity, elongation, convexity, and eccentricity, grouped by eGFP-expression status and vector type, comparing ssAAV2 and scAAV2 to mock. Asterisks indicate significant change relative to mock. * p < 0.05, ** p < 0.01, *** p < 0.001, **** p < 0.0001, ns = not significant.

### Nuclear morphology shifts track the chromatin and polymerase changes

Quantitative nuclear shape metrics are sensitive readouts of chromatin compaction and transcriptional activity, and experimental perturbation of histone modification state is sufficient to alter nuclear rigidity and shape (38). Thus, next we investigated changes in the morphology of the nucleus upon infection. Area, perimeter, convexity, and eccentricity differed from mock in most groups, with absolute Δz typically between 0.1 and 1.0, while circularity and elongation in the self-complementary vector were not significantly different (Fig. 4b).

### ssAAV2 and scAAV2 modulate nuclear protein abundance and LLPS compartment architecture

We quantified the abundance of selected nuclear proteins and morphology of three liquid-liquid phase separated (LLPS) compartments: nucleoli (fibrillarin), promyelocytic leukemia nuclear bodies (PML-bodies, SP100), and nuclear splicing speckles (NSs, SRSF2), at 48 h after infection with ssAAV2 or scAAV2 at a MOI of 20’000 gcp/cell. All measurements are reported as Δz relative to mock to allow direct comparison across features and conditions. Mean nuclear intensities measured over the entire nucleus for fibrillarin, SP100, and SRSF2 were broadly lower for both vectors and both eGFP-expression categories, expressing and non-expressing, compared to mock-infected cells (Fig. 5a). The only group that did not differ from mock was SP100 in eGFP-non-expressing cells upon scAAV2 infection. We then evaluated area, convex area (the area of the convex hull), and solidity of the three selected LLPS compartments. The Δz heatmap in (Fig. 5b) shows condition specific remodeling. In ssAAV2-infected wells, eGFP-non-expressing cells showed decreased PML-body solidity and NSs area, whereas eGFP-expressing cells showed no change for nucleolar solidity or for both, area and solidity of NSs. All other features shifted relative to mock as indicated. Next, we compared the number of compartments per cell, restricting the analysis to cells with at least one compartment. Nucleolar counts were stable across groups for both vectors (Fig. 5c). PML-bodies increased in both eGFP-non-expressing and eGFP-expressing populations, whereas NSs showed a modest increase in number. Representative SRSF2 images (Fig. 5d) illustrate these patterns and provide visual confirmation of the group level effects.

**Fig. 5:**
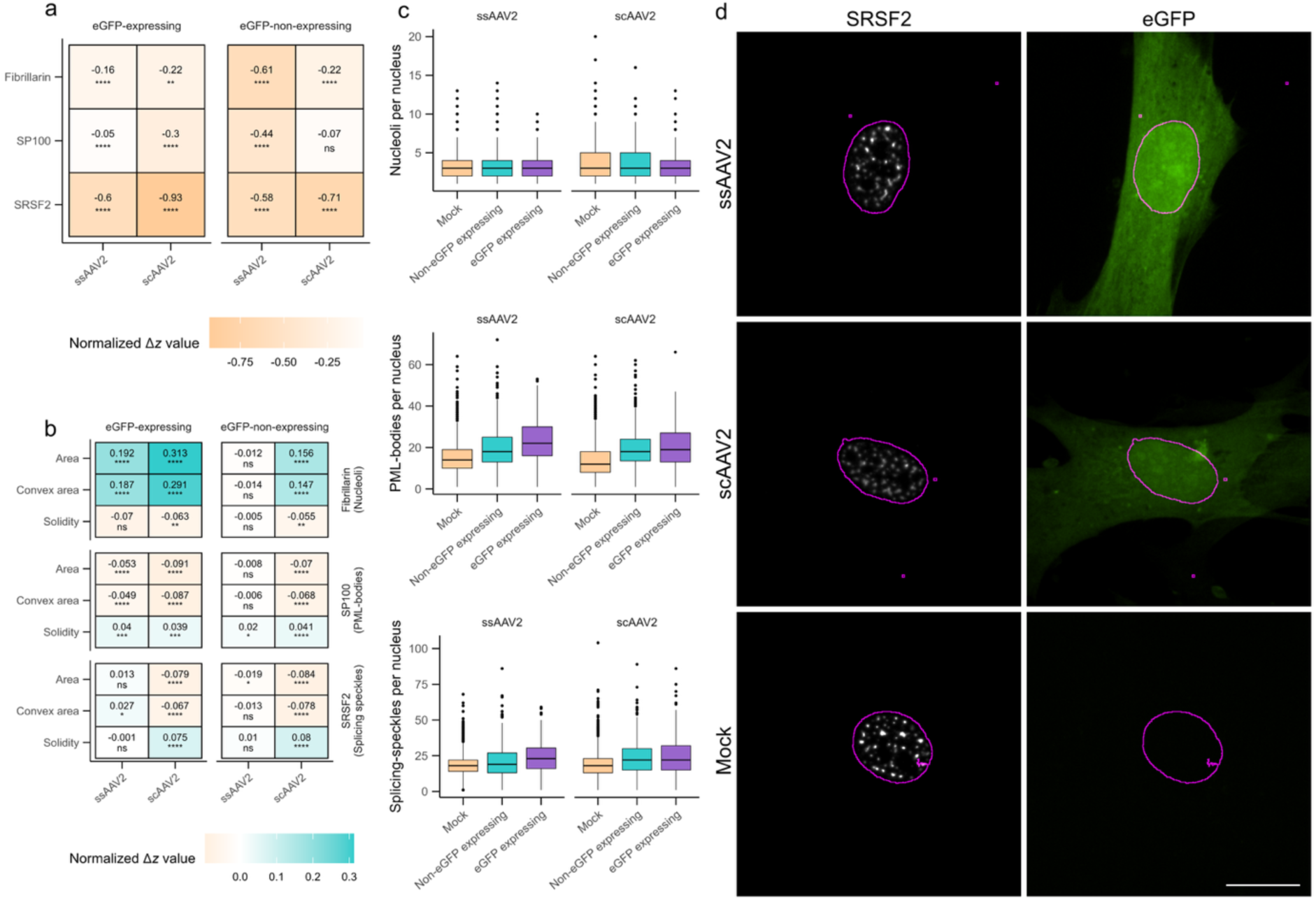
Nuclear protein intensity and LLPS compartment morphology after infection with ssAAV2 and scAAV2. (a) Heatmap of Δz for mean nuclear intensity of fibrillarin, SP100, and SRSF2, grouped by eGFP-non-expressing and eGFP-expressing cells, showing the difference of ssAAV2 or scAAV2 infection cells relative to mock-infected cells. Values were z-score normalized per feature and condition, then Δz computed. All stains are reduced relative to mock, except SP100 in eGFP-non-expressing cells upon scAAV2-infection, which is not significantly changed. (b) Heatmap of Δz for area, convex area, and solidity of nucleoli (fibrillarin), PML-bodies (SP100), and NSs (SRSF2), grouped by eGFP expression and vector type. Asterisks indicate significant change relative to mock. (c) Boxplots of the number of compartments upon infection with ssAAV2 and scAAV2 with groups being mock-infected (mock), eGFP-non-expressing, and eGFP-expressing, restricted to cells with more than zero compartments. (d) Representative images of the SRSF2 stain showing NSs (left) with the eGFP-expression (right) and the nucleus as outline (magenta), scale bar = 10 μm. * p < 0.05, ** p < 0.01, *** p < 0.001, **** p < 0.0001, ns = not significant.

## Discussion

The aim of this study was to evaluate the effect of AAV2 (vector) genomes on the nuclear architecture in primary human fibroblasts. Specifically, we investigated whether entry of wildtype AAV2 or delivery of ssAAV2 or scAAV2 alter host chromatin accessibility using ATAC-seq (Fig. 1). Across two time points, genome-wide ATAC-seq showed no detectable change in host chromatin accessibility, which was confirmed by DNase I digestion assay (Fig. 2) of selected genes (ACTB, GNG7, IFNB1, OSBPL8, and PCNA). This data suggests that at the doses applied, the virus or vectors did not significantly open or close large sets of regulatory elements, as opposed to other viruses such as herpes simplex virus type 1 (HSV-1), which alters chromatin accessibility after just 6 hours (39). The selected MOIs exceed typical *in vivo* doses, hence any patient phenotypes would occur at lower exposure. High dose systemic AAV has been associated with toxicity in animals and humans, including hepatotoxicity and complement-mediated thrombotic microangiopathy (40,41). Even at these higher MOIs we did not detect large scale changes in chromatin accessibility, therefore therapeutic dosing, which is lower, would be expected to yield effects that are at most similar and more likely smaller under comparable conditions.

An alternative explanation is reduced sensitivity to modest changes occurring at dispersed loci. ATAC-seq and DNase I digestion are less effective in detecting subtle alterations that are widespread but shallow, and we cannot rule out focal remodeling that falls below our detection thresholds. Approaches with higher resolution, such as single-cell ATAC-seq, may overcome this limitation. In this context, prior work shows that rAAV vector genomes become chromatinized and accrue histone markers over time *in vitro* and *in vivo*, providing a mechanistic basis for time dependent changes on the vector genome even when host accessibility appears stable (42). Thus, the apparent stability of accessibility should be interpreted as the absence of major effects rather than clear evidence of absence of local change (43).

In contrast to accessibility, nuclear immunofluorescence readouts showed lower mean signals for H3, H3K27me3, and RNA polymerase II markers across groups (Fig. 4). This pattern is consistent with a mild reduction in polymerase engagement and histone mark abundance and matches prior reports that cellular stress and virus infection associate with reduced Ser2 phosphorylation on the RNA polymerase II C-terminal domain (CTD) and altered polymerase turnover, which dampens elongation-linked signals without necessarily shutting down transcription (44,45).

Given that AAV exposure can activate DNA damage signaling, the modest changes in polymerase-associated markers may reflect stress-driven pathways that couple entry to chromatin and transcription, consistent with DNA damage response engagement by AAV and the cell-cycle restricted nature of AAV2 replication (46,47). In line with this, AAV exposure has been reported to induce S or G2 delays in human cells (48).

H3K4me3 deviated from the general trend. While most groups displayed a decrease, G1 cells that were eGFP-expressing upon scAAV2 infection showed an increase in H3K4me3 signal (Fig. 4). One interpretation is selective promoter priming at adaptation genes during a broader dampening of polymerase activity, which has been described in stress models where H3K4me3 can be retained or broadened at specific promoters while declining elsewhere (49,50).

Nuclear geometry changed in parallel with the protein signals. Area, perimeter, convexity, and eccentricity differed from mock in most infected groups, whereas circularity and elongation remained unchanged with the scAAV2 vector (Fig. 4). These metrics are known to respond to chromatin compaction and transcriptional activity, and previous work has shown that experimental modulation of histone modifications can alter nuclear mechanics. In our data, chromatin accessibility remained largely stable, indicating that the modest shape alterations observed are unlikely to arise from broad chromatin remodeling. Instead, they may reflect indirect pathways engaged by AAV exposure, such as stress or DNA damage signaling (51).

Moreover, markers of liquid-liquid phase separated (LLPS) compartments also shifted. Mean total nuclear intensities for fibrillarin (nucleoli), SP100 (PML-bodies), and SRSF2 (NSs) were reduced in transduced cells compared to mock. In addition, structural remodeling was evident, with changes in area, convex area, and solidity, accompanied by increased numbers of PML-bodies and NSs (Fig. 5). Reduced fibrillarin is consistent with nucleolar stress and redistribution of nucleolar components during cellular adaptation to perturbation (52). It may also be linked to AAV2 biology, as the nucleolus has been proposed as a site for uncoating (53).

The number and geometry of PML-bodies are known to respond to interferon and oxidative cues, and combined interferon and DNA damage signaling can generate enlarged compartments that interact with nucleic acids. This aligns with the direction of change we observe, although still within a modest range (54,55). Furthermore, NSs reorganization under stress with recruitment of splicing factors and polymerase has been reported and offers a framework for the SRSF2 findings (56,57).

The implications are both practical and mechanistic. At the tested doses and times, AAV2 and single stranded or self-complementary AAV2 vectors do not appear to remodel host chromatin accessibility at scale, while eliciting mild changes in polymerase-linked signals, nuclear geometry, and condensate architecture.

We used primary human fibroblasts to establish a generalizable concept in a tractable primary cell type, which often also takes the role of a bystander cell, and the framework can be applied to liver, muscle, or other clinically relevant cells. For vector design and safety, these observations support the view that AAV2-based vectors overall induce only modest nuclear alterations while maintaining a favorable safety profile.

## Acknowledgments

We would like to thank H. Büning (Hannover Medical School) for providing purified wildtype AAV2 (wtAAV2). Recombinant AAV2 vectors expressing eGFP (ssAAV2eGFP: p-AME1 and scAAV2eGFP: p-65-2) were supplied by the Viral Vector Facility (VVF) of the Neuroscience Center Zurich (ZNZ). Genomic analyses were performed at the Functional Genomics Center Zurich (FGCZ). The Imaging was performed with equipment maintained by the Center for Microscopy and Image Analysis, University of Zurich.

## Author’s contribution

J.W.Z.: Conceptualization, Methodology, Software, Validation, Formal analysis, Investigation, Data curation, Writing, original draft, Visualization, Project administration. M.K.P.: Conceptualization, Methodology, Software, Validation, Formal analysis, Investigation, Data curation, Writing, original draft, Visualization, Project administration. K.T.: Software, Validation, Formal analysis, Investigation, Data curation, Writing, review and editing, Visualization. C.F.: Conceptualization, Writing, review and editing, Project administration, Supervision, Funding acquisition.

## Supplementary material

Supplementary Tab. 1

Supplementary Tab. 2

Supplementary Tab. 3

## References

1. Au HKE, Isalan M, Mielcarek M. Gene Therapy Advances: A Meta-Analysis of AAV Usage in Clinical Settings. Front Med [Internet]. 2022 [cited 2023 Mar 29];8. Available from: https://www.frontiersin.org/articles/10.3389/fmed.2021.809118

2. Burdett T, Nuseibeh S. Changing trends in the development of AAV-based gene therapies: a meta-analysis of past and present therapies. Gene Ther. 2023 Apr;30(3):323–35.

3. Pipe SW, Leebeek FWG, Recht M, Key NS, Castaman G, Miesbach W, et al. Gene Therapy with Etranacogene Dezaparvovec for Hemophilia B. N Engl J Med. 2023 Feb 23;388(8):706–18.

4. Zwi-Dantsis L, Mohamed S, Massaro G, Moeendarbary E. Adeno-Associated Virus Vectors: Principles, Practices, and Prospects in Gene Therapy. Viruses. 2025 Feb 9;17(2):239.

5. Samulski RJ, Muzyczka N. AAV-Mediated Gene Therapy for Research and Therapeutic Purposes. Annu Rev Virol. 2014;1(1):427–51.

6. Gil-Farina I, Fronza R, Kaeppel C, Lopez-Franco E, Ferreira V, D’Avola D, et al. Recombinant AAV Integration Is Not Associated With Hepatic Genotoxicity in Nonhuman Primates and Patients. Mol Ther J Am Soc Gene Ther. 2016 June;24(6):1100–5.

7. Kaeppel C, Beattie SG, Fronza R, van Logtenstein R, Salmon F, Schmidt S, et al. A largely random AAV integration profile after LPLD gene therapy. Nat Med. 2013 July;19(7):889–91.

8. McCarty DM, Young SM, Samulski RJ. Integration of Adeno-Associated Virus (AAV) and Recombinant AAV Vectors. Annu Rev Genet. 2004;38(1):819–45.

9. Nakai H, Yant SR, Storm TA, Fuess S, Meuse L, Kay MA. Extrachromosomal Recombinant Adeno-Associated Virus Vector Genomes Are Primarily Responsible for Stable Liver Transduction In Vivo. J Virol. 2001 Aug;75(15):6969.

10. Schnepp BC, Jensen RL, Chen CL, Johnson PR, Clark KR. Characterization of Adeno-Associated Virus Genomes Isolated from Human Tissues. J Virol. 2005 Dec 15;79(23):14793–803.

11. Penaud-Budloo M, Le Guiner C, Nowrouzi A, Toromanoff A, Chérel Y, Chenuaud P, et al. Adeno-Associated Virus Vector Genomes Persist as Episomal Chromatin in Primate Muscle. J Virol. 2008 Aug;82(16):7875–85.

12. Westhaus A, Cabanes-Creus M, Rybicki A, Baltazar G, Navarro RG, Zhu E, et al. High-Throughput In Vitro, Ex Vivo, and In Vivo Screen of Adeno-Associated Virus Vectors Based on Physical and Functional Transduction. Hum Gene Ther. 2020 May;31(9–10):575–89.

13. Schnepp BC, Chulay JD, Ye GJ, Flotte TR, Trapnell BC, Johnson PR. Recombinant Adeno-Associated Virus Vector Genomes Take the Form of Long-Lived, Transcriptionally Competent Episomes in Human Muscle. Hum Gene Ther. 2016 Jan;27(1):32–42.

14. Kong X, Wei G, Chen N, Zhao S, Shen Y, Zhang J, et al. Dynamic chromatin accessibility profiling reveals changes in host genome organization in response to baculovirus infection. PLOS Pathog. 2020 June 8;16(6):e1008633.

15. Callé A, Ugrinova I, Epstein AL, Bouvet P, Diaz JJ, Greco A. Nucleolin is required for an efficient herpes simplex virus type 1 infection. J Virol. 2008 May;82(10):4762–73.

16. Gu H, Roizman B. The degradation of promyelocytic leukemia and Sp100 proteins by herpes simplex virus 1 is mediated by the ubiquitin-conjugating enzyme UbcH5a. Proc Natl Acad Sci. 2003 July 22;100(15):8963–8.

17. Wingett SW, Andrews S. FastQ Screen: A tool for multi-genome mapping and quality control. F1000Research. 2018 Sept 17;7:1338.

18. Andrews S. Babraham Bioinformatics - FastQC A Quality Control tool for High Throughput Sequence Data [Internet]. 2010 [cited 2025 Aug 22]. Available from: https://www.bioinformatics.babraham.ac.uk/projects/fastqc/

19. Langmead B, Salzberg SL. Fast gapped-read alignment with Bowtie 2. Nat Methods. 2012 Apr;9(4):357–9.

20. Hatakeyama M, Opitz L, Russo G, Qi W, Schlapbach R, Rehrauer H. SUSHI: an exquisite recipe for fully documented, reproducible and reusable NGS data analysis. BMC Bioinformatics. 2016 June 2;17(1):228.

21. Amemiya HM, Kundaje A, Boyle AP. The ENCODE Blacklist: Identification of Problematic Regions of the Genome. Sci Rep. 2019 June 27;9(1):9354.

22. Ramírez F, Ryan DP, Grüning B, Bhardwaj V, Kilpert F, Richter AS, et al. deepTools2: a next generation web server for deep-sequencing data analysis. Nucleic Acids Res. 2016 July 8;44(W1):W160–5.

23. Zhang Y, Liu T, Meyer CA, Eeckhoute J, Johnson DS, Bernstein BE, et al. Model-based Analysis of ChIP-Seq (MACS). Genome Biol. 2008 Sept 17;9(9):R137.

24. Quinlan AR, Hall IM. BEDTools: a flexible suite of utilities for comparing genomic features. Bioinformatics. 2010 Mar 15;26(6):841–2.

25. Li H, Handsaker B, Wysoker A, Fennell T, Ruan J, Homer N, et al. The Sequence Alignment/Map format and SAMtools. Bioinformatics. 2009 Aug 15;25(16):2078–9.

26. Ewels P, Magnusson M, Lundin S, Käller M. MultiQC: summarize analysis results for multiple tools and samples in a single report. Bioinformatics. 2016 Oct 1;32(19):3047–8.

27. Stark R, Brown GD. DiffBind : Differential binding analysis of ChIP-Seq peak data. In 2012. Available from: https://api.semanticscholar.org/CorpusID:1875957

28. Wickham H. ggplot2: Elegant Graphics for Data Analysis. 2nd ed. 2016. Cham: Springer International Publishing : Imprint: Springer; 2016. 1 p. (Use R!).

29. Wickham H, Averick M, Bryan J, Chang W, McGowan L, François R, et al. Welcome to the Tidyverse. J Open Source Softw. 2019 Nov 21;4(43):1686.

30. Sofroniew N, Lambert T, Evans K, Nunez-Iglesias J, Bokota G, Winston P, et al. napari: a multi-dimensional image viewer for Python [Internet]. Zenodo; 2022 [cited 2025 Aug 22]. Available from: https://zenodo.org/record/7276432

31. Schindelin J, Arganda-Carreras I, Frise E, Kaynig V, Longair M, Pietzsch T, et al. Fiji: an open-source platform for biological-image analysis. Nat Methods. 2012 July;9(7):676–82.

32. Rämö P, Sacher R, Snijder B, Begemann B, Pelkmans L. CellClassifier: supervised learning of cellular phenotypes. Bioinforma Oxf Engl. 2009 Nov 15;25(22):3028–30.

33. R: The R Project for Statistical Computing [Internet]. [cited 2023 Mar 8]. Available from: https://www.r-project.org/

34. Posit team. RStudio: Integrated Development Environment for R [Internet]. Boston, MA: Posit Software, PBC; 2025. Available from: http://www.posit.co/

35. Nepon-Sixt B, Alexandrow M. DNase I Chromatin Accessibility Analysis. BIO-Protoc [Internet]. 2019 [cited 2024 Nov 12];9(23). Available from: https://bio-protocol.org/e3444

36. Fetni R, Richer CL, Malfoy B, Dutrillaux B, Lemieux N. Cytologic characterization of two distinct alpha satellite DNA domains on human chromosome 7, using double-labeling hybridizations in fluorescence and electron microscopy on a melanoma cell line. Cancer Genet Cytogenet. 1997 July;96(1):17–22.

37. Narotamo H, Fernandes MS, Moreira AM, Melo S, Seruca R, Silveira M, et al. A machine learning approach for single cell interphase cell cycle staging. Sci Rep. 2021 Sept 29;11(1):19278.

38. Stephens AD, Liu PZ, Banigan EJ, Almassalha LM, Backman V, Adam SA, et al. Chromatin histone modifications and rigidity affect nuclear morphology independent of lamins. Misteli T, editor. Mol Biol Cell. 2018 Jan 15;29(2):220–33.

39. Hennig T, Michalski M, Rutkowski AJ, Djakovic L, Whisnant AW, Friedl MS, et al. HSV-1-induced disruption of transcription termination resembles a cellular stress response but selectively increases chromatin accessibility downstream of genes. Kalejta RF, editor. PLOS Pathog. 2018 Mar 26;14(3):e1006954.

40. Hinderer C, Katz N, Buza EL, Dyer C, Goode T, Bell P, et al. Severe Toxicity in Nonhuman Primates and Piglets Following High-Dose Intravenous Administration of an Adeno-Associated Virus Vector Expressing Human SMN. Hum Gene Ther. 2018 Mar;29(3):285–98.

41. Hordeaux J, Lamontagne RJ, Song C, Buchlis G, Dyer C, Buza EL, et al. High-dose systemic adeno-associated virus vector administration causes liver and sinusoidal endothelial cell injury. Mol Ther. 2024 Apr;32(4):952–68.

42. Maurer AC, Benyamini B, Whitney ON, Fan VB, Cattoglio C, Kissiov DU, et al. Double-strand break repair pathways differentially affect processing and transduction by dual AAV vectors. Nat Commun. 2025 Feb 11;16(1):1532.

43. Kiani K, Sanford EM, Goyal Y, Raj A. Changes in chromatin accessibility are not concordant with transcriptional changes for single-factor perturbations. Mol Syst Biol. 2022 Sept;18(9):e10979.

44. Yamazaki T, Liu L, Manley JL. Oxidative stress induces Ser 2 dephosphorylation of the RNA polymerase II CTD and premature transcription termination. Transcription. 2021 Oct 20;12(5):277–93.

45. Gulyas L, Glaunsinger BA. RNA polymerase II subunit modulation during viral infection and cellular stress. Curr Opin Virol. 2022 Oct;56:101259.

46. Schwartz RA, Carson CT, Schuberth C, Weitzman MD. Adeno-Associated Virus Replication Induces a DNA Damage Response Coordinated by DNA-Dependent Protein Kinase. J Virol. 2009 June 15;83(12):6269–78.

47. Franzoso FD, Seyffert M, Vogel R, Yakimovich A, De Andrade Pereira B, Meier AF, et al. Cell Cycle-Dependent Expression of Adeno-Associated Virus 2 (AAV2) Rep in Coinfections with Herpes Simplex Virus 1 (HSV-1) Gives Rise to a Mosaic of Cells Replicating either AAV2 or HSV-1. Longnecker RM, editor. J Virol. 2017 Aug;91(15):e00357–17.

48. Zubkova ES, Beloglazova IB, Ratner EI, Dyikanov DT, Dergilev KV, Menshikov MYu, et al. Transduction of rat and human adipose-tissue derived mesenchymal stromal cells by adeno-associated viral vector serotype DJ. Biol Open. 2021 Sept 15;10(9):bio058461.

49. Batie M, Frost J, Frost M, Wilson JW, Schofield P, Rocha S. Hypoxia induces rapid changes to histone methylation and reprograms chromatin. Science. 2019 Mar 15;363(6432):1222–6.

50. Rytkönen KT, Faux T, Mahmoudian M, Heinosalo T, Nnamani MC, Perheentupa A, et al. Histone H3K4me3 breadth in hypoxia reveals endometrial core functions and stress adaptation linked to endometriosis. iScience. 2022 May;25(5):104235.

51. Stephens AD, Liu PZ, Banigan EJ, Almassalha LM, Backman V, Adam SA, et al. Chromatin histone modifications and rigidity affect nuclear morphology independent of lamins. Misteli T, editor. Mol Biol Cell. 2018 Jan 15;29(2):220–33.

52. Boulon S, Westman BJ, Hutten S, Boisvert FM, Lamond AI. The Nucleolus under Stress. Mol Cell. 2010 Oct;40(2):216–27.

53. Sutter SO, Lkharrazi A, Schraner EM, Michaelsen K, Meier AF, Marx J, et al. Adeno-associated virus type 2 (AAV2) uncoating is a stepwise process and is linked to structural reorganization of the nucleolus. Weitzman MD, editor. PLOS Pathog. 2022 July 11;18(7):e1010187.

54. Sahin U, Ferhi O, Jeanne M, Benhenda S, Berthier C, Jollivet F, et al. Oxidative stress–induced assembly of PML nuclear bodies controls sumoylation of partner proteins. J Cell Biol. 2014 Mar 17;204(6):931–45.

55. Scherer M, Read C, Neusser G, Kranz C, Kuderna AK, Müller R, et al. Dual signaling via interferon and DNA damage response elicits entrapment by giant PML nuclear bodies. eLife. 2022 Mar 23;11:e73006.

56. Galganski L, Urbanek MO, Krzyzosiak WJ. Nuclear speckles: molecular organization, biological function and role in disease. Nucleic Acids Res. 2017 Oct 13;45(18):10350–68.

57. Sung HM, Schott J, Boss P, Lehmann JA, Hardt MR, Lindner D, et al. Stress-induced nuclear speckle reorganization is linked to activation of immediate early gene splicing. J Cell Biol. 2023 Dec 4;222(12):e202111151.

