## SupplementaryTables for "The limited effects of AAV2 vectors on host chromatin accessibility and nuclear architecture are consistent with a favorable safety profile"

**Supplementary tables**

Supplementary Tab. 1: qPCR primers used for genome copy determination.

| **Name of the primer** | **Sequence (5' -> 3')** |
| --- | --- |
| p-AV2 FW | ATT GAC GGG AAC TCA ACG AC |
| p-AV2 REV | ATT CAT GCT CCA CCT CAA CC |
| GFP FW | ATT GAC GGG AAC TCA ACG AC |
| GFP REV | ATT CAT GCT CCA CCT CAA CC |
| GAPDH FW | TGC ACC ACC AAC TGC TTA GC |
| GAPDH REV | GGC ATG GAC TGT GGT CAT GAG |

Supplementary Tab. 2: Antibodies used in the HCS-IF.

| **Target** | **Cat.-Nr.** | **Manufacturer** | **Dilution** |
| --- | --- | --- | --- |
| H3 | ab1791 | Abcam | 1:1000 |
| H3K4me3 | ab8580 | Abcam | 1:800 |
| H3K27me3 | 39055 | Active Motif | 1:250 |
| RNAP II | 05-623-Z | Sigma-Aldrich | 1:400 |
| RNA Pol II p-Ser2 | 04-1571 | Sigma-Aldrich | 1:500 |
| RNA Pol II p-Ser5 | ab5408 | Abcam | 1:1500 |
| RNA Pol II p-Ser7 | ab252853 | Abcam | 1:100 |
| Fibrillarin | ab5821 | Abcam | 1:500 |
| SP100 | HPA017384 | Sigma-Aldrich | 1:800 |
| SRSF2 | S4045 | Sigma-Aldrich | 1:1200 |
| Donkey anti-Rat IgG (H+L) Highly Cross-Adsorbed Secondary Antibody, Alexa Fluor™ 568 | A78946 | Thermo Fisher | 1:500 |
| Donkey anti-Mouse IgG (H+L) Highly Cross-Adsorbed Secondary Antibody, Alexa Fluor™ 568 | A10037 | Thermo Fisher | 1:500 |
| Donkey anti-Rabbit IgG (H+L) Highly Cross-Adsorbed Secondary Antibody, Alexa Fluor™ Plus 647 | A32795 | Thermo Fisher | 1:500 |

Supplementary Tab. 3: qPCR primers for DNase I digestion assay.

| **Name of the primer** | **Sequence (5' -> 3')** |
| --- | --- |
| GNG7_FW | AAT GGT GCT GCT GTG AAC CCG G |
| GNG7_REV | TCG TTC CGG GCA TGT TGC TCA C |
| OSBPL8_FW | TGC CTT AGC GGT AGT ATC CTG GC |
| OSBPL8_REV | CCT CCC AGG AGA GGG TGC AAG T |
| ACTB_FW | GCT CTT GCC AAT GGG GAT CGC A |
| ACTB_REV | gca gtt agc gcc caa agg acc a |
| D7Z1_FW | acc gca ttg agg cct tcg ttg g |
| D7Z1_REV | ccg ctc tgt gca aag gga cgt t |
| IFNB1_FW | ACA GTC ACT GTG CCT GGA CCA T |
| IFNB1_REV | TGT CCA GTC CCA GAG GCA CAG G |
| PCNA_FW | TGC AGA GCA TGG ACT CGT CCC A |
| PCNA_REV | GCT CAC CTG GTG AGG TTC ACG C |
